# Degenerate time-dependent network dynamics anticipate seizures in human epileptic brain

**DOI:** 10.1101/080739

**Authors:** Adrià Tauste Campo, Alessandro Principe, Miguel Ley, Rodrigo Rocamora, Gustavo Deco

## Abstract

Epileptic seizures are known to follow specific changes in brain dynamics. While some algorithms can nowadays robustly detect these changes, a clear understanding of the mechanism by which these alterations occur and generate seizures is still lacking. Here, we provide cross-validated evidence that such changes are initiated by an alteration of physiological network state dynamics. Specifically, our analysis of long intracranial EEG recordings from a group of 10 patients identifies a critical phase of a few hours in which time-dependent network states become less variable (“degenerate”) and is followed by a global functional connectivity reduction before seizure onset. This critical phase is characterized by an abnormal occurrence of highly correlated network instances and is shown to particularly affect the activity of resection regions in patients with validated postsurgical outcome. Our approach characterizes pre-seizure networks dynamics as a cascade of two sequential events providing new insights into seizure prediction and control.

## Introduction

Epilepsy is among the most common neurological disorders with an estimated prevalence of about 1% of the world’s population and almost 2% in low-income families in developed countries (CDC, 2010). Epilepsy is characterized by the seemingly random occurrence of seizures, which can greatly affect the quality of life of patients. Approximately one third of all epileptic patients are resistant to pharmacotherapy (Kwan *et al*., 2011) and could benefit from a variety of surgical options. Among them, closed-loop neuromodulation based on an accurate prediction of seizure occurrences is a promising tool.

Over the last decades, several studies have showed that seizures are preceded by detectable changes in brain dynamics that can be measured via intracranial recordings. Although not being fully understood, these changes have been associated to the existence of a transition from interictal activity to *pre-ictal state* (Lopes da Silva 2003, Stacey *et al*., 2011). These findings have motivated intense research on the development of seizure prediction algorithms for therapeutic use in patients with refractory epilepsy (Park *et al*., 2011, Valderrama *et al*., 2012, Cook *et al*., 2013, Gadhoumi *et al*., 2015). Although significant progress has been made to attain above-chance level performance results (Brinkmann *et al*., 2016), there is yet a long road to turn seizure prediction into therapeutic devices (Freestone *et al*., 2017). A major caveat of current seizure prediction is the lack of understanding about the neurophysiological processes associated to the emergence and maintenance of the pre-ictal state. Indeed, most studies have resorted to fully data-driven methods to discriminate the pre-ictal state with multiple signal features, which are typically patient-specific and difficult to interpret (Gadhoumi *et al*., 2015).

Nowadays, epilepsy research is gradually adopting a network approach to study seizure dynamics at a global level and assess the contribution of the epileptogenic zone (Van Diessen *et al*. 2013, Van Mierlo *et al*., 2014, Goodfellow *et al*., 2016, Khambati *et al*., 2016). In this growing field, the majority of published studies have identified specific graph-theoretical properties of functional networks during ictal and interictal periods (Kramer *et al*., 2008, Bartolomei *et al*., 2011, Haneef *et al*., 2014, Stam 2014). In particular, a few groups have started to characterize the temporal variability of such functional networks during ictal (Rummel *et al*. 2013, Burns *et al*., 2014, Khambati *et al*., 2015) and interictal epochs (Takahashi *et al*., 2012, Geier *et al*., 2015a, Khambati *et al*., 2017). Specifically, some authors have employed state spaces to classify recurrent functional networks during seizures to pinpoint those states that were responsible for the generation, maintenance and termination of ictal activity (Burns *et al*., 2014, Khambati *et al*., 2015). More recently, a similar approach has been applied to a large sample of 10-minutes interictal epochs showing that interictal activity exhibits larger fluctuations than ictal periods over a common set of states (Khambati *et al*., 2017). In this context, however, the crucial question on whether there exist network dynamics changes pointing towards an upcoming seizure remains unaddressed. It is therefore due to ask: (1) how are recurrent network states dynamically altered before epileptic seizures? More generally, can network dynamics provide a common principle of the pre-ictal state?

In the current study, we addressed these questions for the first time by analyzing time-dependent alterations in the dynamic repertoire of the functional connectivity (Hutchinson *et al*., 2013) during long pre-seizure periods preceding seizures. Based on insights from other dysfunctional models (Hudetz *et al*., 2014, Barttfeld *et al*., 2015) and recent findings showing network dynamics alterations between interictal and ictal epochs (Khambati *et al*., 2017), we hypothesized that the variability of physiological (non-dysfunctional) network states was reduced as interictal activity approached epileptic seizures. Under this hypothesis, we developed a novel analysis to study specific variability changes prior to seizures preceded by long interictal periods in 10 epileptic patients monitored with video-SEEG (stereoencephalography) during pre-surgical diagnosis. We made use of a graph-theoretical property, the eigenvector centrality, to characterize network states (Burns *et al*., 2014) as instances of a time-varying multivariate continuous variable, and resorted to the Gaussian entropy (Cover and Thomas, 2012) to describe their variability. A controlled analysis using time-matched periods of interictal activity from additional days revealed a consistent and sustained decrease of the variability of network states before the seizure occurred. Remarkably, in all patients this loss of variability was specifically associated to an abnormal occurrence of high-connectivity states during the pre-seizure period. We also investigated the contribution of the epileptogenic sites to the measured effect in two patients with a long-lasting (>4 years) very good post-operative outcome. In particular, the application of our analysis to the mapped epileptogenic sites of these seizure-free patients showed a significant alteration in the resected areas of the patients’ epileptic networks. Overall, our approach provides two main contributions in the analysis of epileptic network dynamics. First, it characterizes the pre-ictal state as a two-stage process in which epileptic networks undergo a functional reorganization before seizure onset. Second, it develops methodological aspects that may be considered to improve seizure prediction algorithms. More broadly, the results presented here open new lines to investigate critical alterations in pathological networks by studying the time-varying nature of brain networks.

## Results

We studied network dynamics prior to epileptic seizures in 10 drug-resistant patients using continuous multichannel intracranial recordings via stereoelectroencephalography (SEEG) during pre-surgical monitoring evaluation (See details in Fig 1). To capture long-term changes in network dynamics, we considered patients whose first spontaneous clinical seizure occurred after at least 30 hours (average value: 71.4±19.1 hours; mean±std) of intracranial implantation. This ictal activity exhibited variable onset times over patients that were more concentrated during the 0:00-8:00 period (Fig 2A). For every patient, we analyzed a long continuous period (average value: 10.4±1.9 hours; mean±std) of intracranial activity before the seizure occurred (pre-seizure period, Fig 2B). We controlled for the specificity of our findings by independently analyzing time-matched periods of interictal activity from different days (e.g., control period, Fig 2B). In this study, we separately analyzed eight patients (Patients 1-8, Main patients) with no clinically relevant events before the first seizure and two patients that presented potential factors perturbing the pre-seizure period (Patients 9 and 10, Control patients). More precisely, Patient 9 had been electrically stimulated 16.5 hours before the first recorded seizure and Patient 10 presented a subclinical seizure 6.1 hours before the first clinical seizure onset.

**Fig 1.**
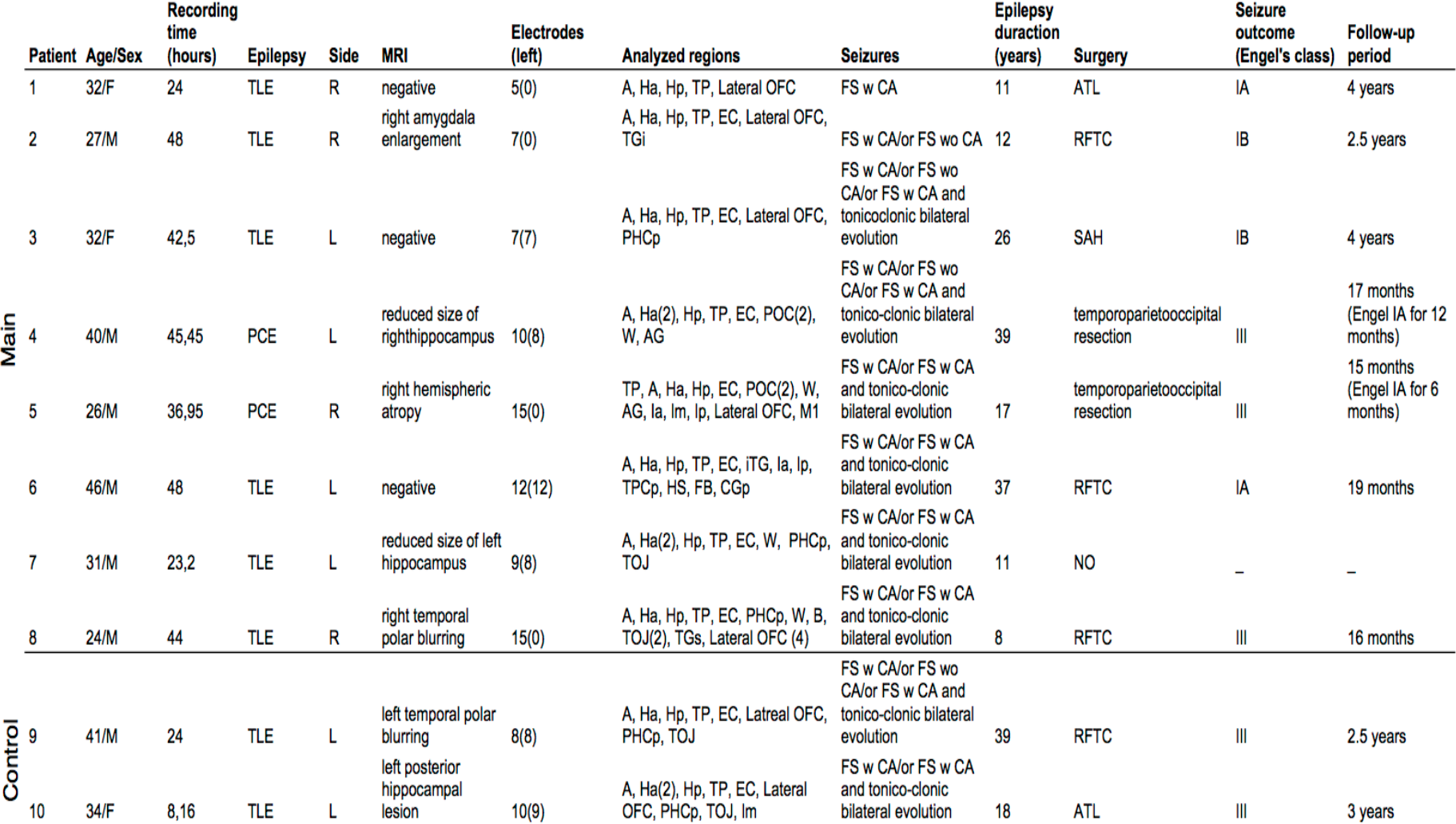
Main data of patients included in the study. F = female; M = male; TLE=temporal lobe epilepsy; PCE=posterior cortex epilepsy; R=right; L=left; A=amygdala; Ha=anterior hippocampus; Hp=posterior hippocampus; TP=temporal pole; EC=entorhinal cortex, Lateral OFC=lateral parts of the orbitofrontal cortex; TGi=inferior temporal gyrus; PHCp=posterior parahippocampal cortex; W=Wernicke’s area; AG=angular gyrus; Ia=anterior insula; Im=mid insula; Ip=posterior insula; M1=primary motor area; TPCp=posterior temporoparietal cortex; HS=Heschl’s area; FB=frontobasal area; CGp=posterior cingulate; TGs=superior temporal gyrus; TOJ=temporal occipital junction; POC=precuneus occipital cortex; B=Broca’s area; FS=focal seizure; w=with; wo=without; CA=consciousness alteration; ATL: Anterior temporal lobectomy; RFTC=Radiofrequency thermocoagulation; SAH=Selective amygdalohyppocampectomy; NO=not-operated.

**Fig 2.**
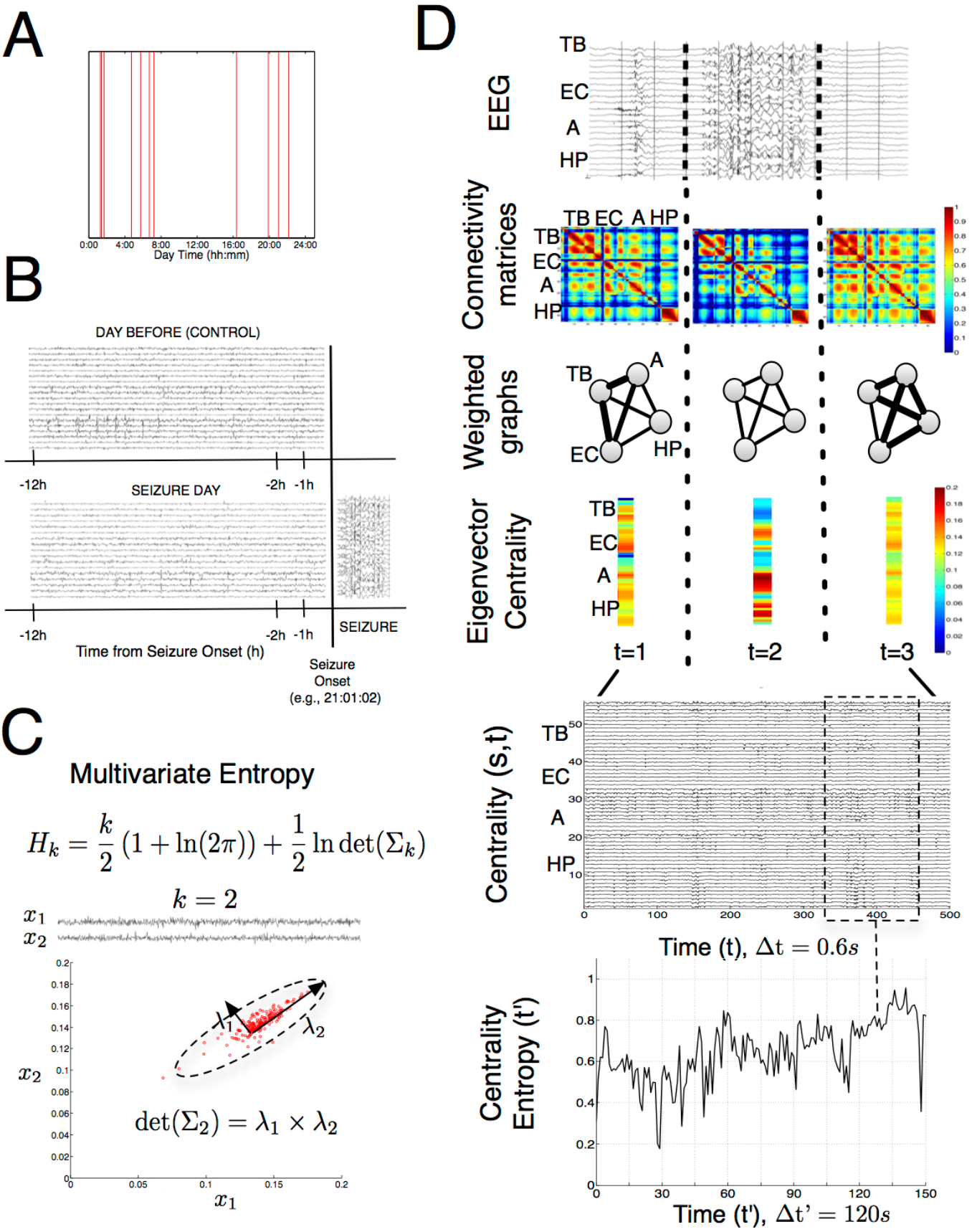
Study paradigm and network dynamics analysis. (A) Seizure onset time of the first recorded spontaneous clinical seizure from every patient (n=10). (B) Schematic representation of the experimental design: for each patient a pre-seizure period of up to 12 hours was matched to the same time period of the previous day that served as a baseline reference (control interictal period). (C) Multivariate (Gaussian) entropy, showing its dependence on the determinant of the covariance matrix (). Example for a case of two time series in which the determinant of the covariance is shown to shape the joint variability. (D) Network dynamics analysis: Simultaneous EEG recordings were first divided into consecutive and non-overlapping time windows of 0.6s (Top). Then, functional connectivity matrices were computed using zero-lagged absolute-valued Pearson correlation in each time windows (Middle-top 1). These matrices were modeled as weighted undirected graphs where nodes represented recorded contacts and edges strength represented correlation absolute values (Middle-top 2). The centrality of each contact in every graph was evaluated using the eigenvector centrality leading to a sequence of centrality vectors (Middle-bottom 1). The overall eigenvector centrality sequence was regarded as a set of simultaneous centrality time series (one for each patient recording site over time steps of 0.6s (Middle-bottom 2). Finally, time-dependent centrality entropy values were found for each period of interest by sequentially estimating the multivariate entropy of the centrality time series in non-overlapping and consecutive time windows of 120s (200 samples). The labels TB (Temporal basal area), EC (Entorhinal cortex), A (Amygdala), and HP (Hippocampus) are used as an example to illustrate where the anatomical information was conveyed in each step of the analysis

### Network dynamics analysis

We tracked network state dynamics for each patient separately over each SEEG recording session. To do so, we computed functional connectivity using Pearson correlation across all recording sites (also referred to as sites, average value: 98.3±25.1 sites; mean±std) over consecutive and non-overlapping time windows of 0.6s (Fig 2D). Networks in each window were characterized as a weighted undirected graph, where electrode contacts represented the nodes and absolute-valued pairwise correlations represented their weighted edges (Fig 2D). We then evaluated a centrality measure for each connectivity matrix to track network dynamics in a reduced and interpretable dimensionality space. Indeed, we computed the eigenvector centrality to reduce each *N x N* connectivity matrix to a N-dimensional vector, where *N* was the total number of recording sites, thus obtaining a centrality sequence for each recording site (Fig 2D). This measure can be equivalently interpreted as the first principal component of the normalized covariance matrix of the set of intracranial recordings in each window.

Our initial hypothesis was that the pre-ictal state was associated with a reduction of physiological network states. We therefore tested this hypothesis by quantifying changes in the distribution of the eigenvector centrality sequences representing these network states. In particular, we assumed that the centrality time series could be approximated by a multivariate Gaussian distribution for a sufficiently large number of samples (n>100). In principle, the second-order variability of a multivariate variable may exhibit two components: the temporal component, i.e., how the centrality of a recording site varies as a function of time, and the spatial component, i.e., how the centrality consistently varies across recording sites at a given time instance. A measure that simultaneously quantifies both components is the multivariate Gaussian entropy, which monotonically depends on the product of the covariance matrix’s eigenvalues (Fig 2C). This measure corresponds to the differential entropy of multivariate normally distributed variables (Cover and Thomas, 2012) but it can be proved useful to approximate the variability of more general variables whose distribution is asymptotically Gaussian (Chen *et al*., 2010).

### Network state variability identifies time-dependent alterations before seizure onset

First, we centered our analysis on the pre-seizure period and the time-matched period from the previous day (pre-seizure, control). Over both periods we computed the multivariate Gaussian entropy in consecutive and non-overlapping time windows of 200 centrality samples (120s) and normalized the measure to lie within the interval [0,1] per patient. We shall refer to this applied measure as centrality entropy in the remaining of the article. The straightforward application of the centrality entropy to both periods in the main patients showed that centrality sequences were generally less entropic during the pre-seizure period (See Fig S1A) showing a gradual increase and successive decrease of this cross-period difference as the seizure onset approached. In order to localize this effect in a specific and significant time segment, we grouped consecutive entropy values into intervals and made use of a non-parametric test to identify the cluster of consecutive centrality entropy intervals that was significantly yielding the largest entropy decay per patient (*Materials and Methods*). The results of this test are illustrated for the main patients in Fig 3A where average centrality entropy curves are plotted for the control (in blue) and pre-seizure period (in red) together with the identified significant time segment (in cyan) during the 9.5 hours preceding the seizure. In each patient, this segment highlighted intervals where the same centrality entropy reduction could not be achieved by shuffling the entropy values within each interval across the pre-seizure and control periods (P<0.01, Fig S1B). Intriguingly, the pinpointed segment was rather patient-specific exhibiting offset times that were not generally attached to the seizure onset. However, when grouping samples across the main patients, significant intervals turned out to be regularly distributed around the proximity of the seizure onset with the interval [-2.5, -1.5] being the most frequent (87.5%, Fig 3B). In particular, this distribution was statistically different (P<0.01, Kolmogorov-Smirnov test) from a surrogate distribution obtained by randomly placing the same segments per patient in every possible location of the pre-seizure period (Fig 3C). In addition, relevant features of the significant segment such as the onset and offset times, and the test’s statistic value were not correlated with the seizure onset time (Fig S2B, C and D). These findings corroborated that our analysis controlled for posible underlying circadian modulations of the iEEG data (Fig S2A). Finally, the results obtained in both control patients were rather different between each other (Fig S3A). In particular, the cross-period difference measured in Patient 9 was the least significant across all patients (Fig S3B), suggesting that the previous received electrical stimulation might have had an effect on the pre-seizure dynamics. In contrast, the occurrence of a subclinical seizure in Patient 10 did not yield a quantitatively different significance effect. We analyzed the stability of the results over the main patients using a synchronization measure (phase-locking value) over difference frequency bands and an alternative network measure (node strength, *Materials and Methods*). The separate application of both measures unravelled similar trends with weaker statistical effects (Figs. S4 and S5). In conclusion, our initial findings suggested that significant and sustained reductions of network state variability over a precedent-day baseline could be related to a pre-ictal state. Further, this reduction in variability was statistically mapped to a patient-specific time sub-period per patient. This sub-period will be referred to in the following as the critical phase.

**Fig 3.**
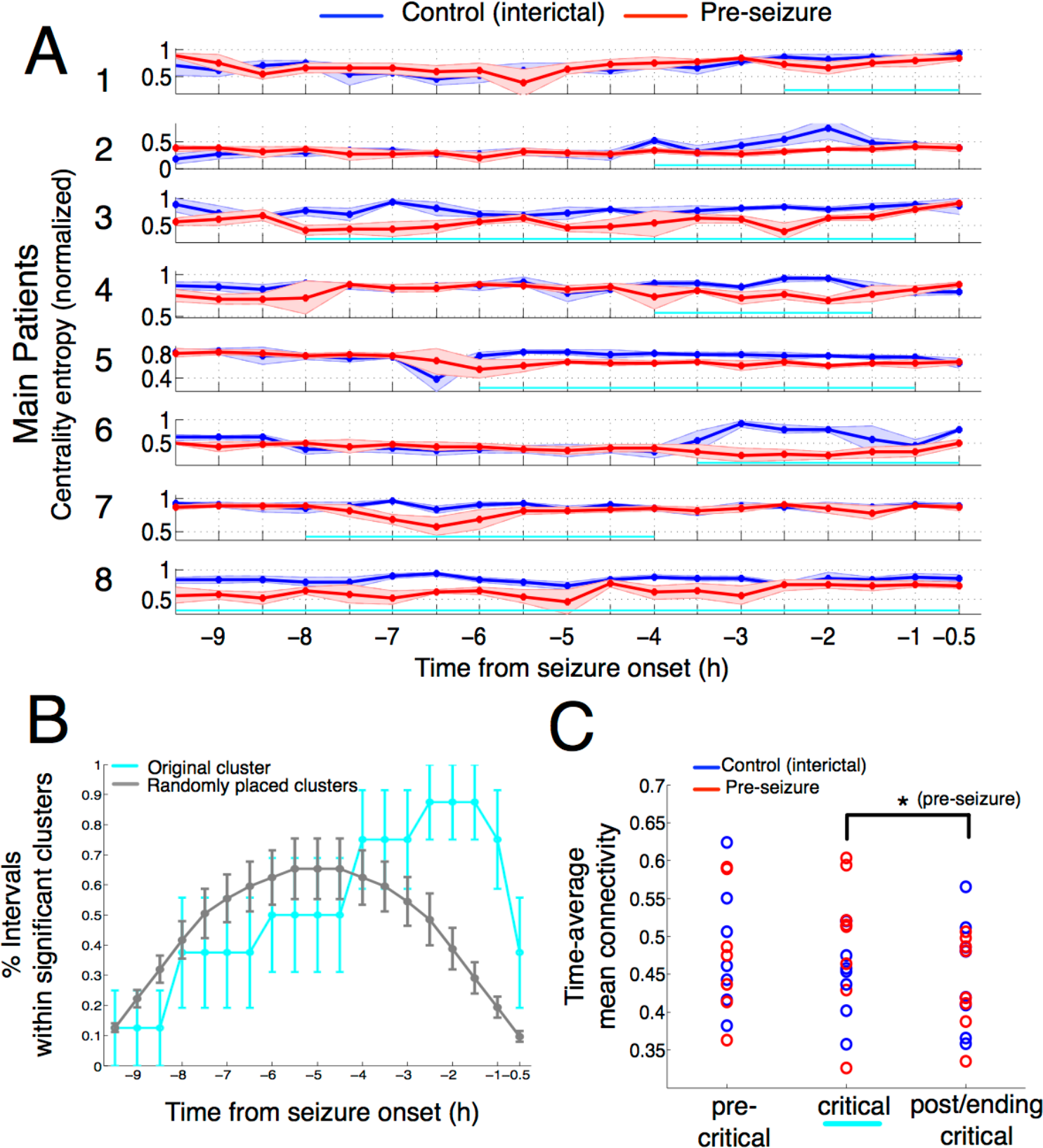
Time-dependent network state variability decreases near seizure onset during pre-seizure periods. (A) Average normalized (to the [0,1] range) centrality entropy for the main epileptic patients (n=8) during a preseizure period (in red, 9.5 hours before the first seizure) and a control period (in blue, 9.5 hours from the preceding day). Averages were computed over time in non-overlapping windows of 15 entropy samples each (total of 30 min) during both periods. Each entropy sample was computed in a smaller window of 200 subsamples (120s). Curves represent the sequence of centrality entropy mean values and error bars represent ± one standard deviation. In cyan, the sequence of consecutive time steps lying in a significant clusterized difference (randomization test, P<0.01). (B) Percentage of times that 30-minute intervals lie within a significant cluster. In cyan, significant clusters are located in their original position. In grey, significant clusters are randomly placed along the pre-seizure periods of each patient. Error bars represent ± SEM (standard error of the mean). (C) Median (across patients) of the time-average mean functional connectivity along three consecutive sub-periods of interest during pre-seizure and control periods. The first sub-period (pre-critical) comprises intervals prior to the significant cluster, the intermediate sub-period (critical) comprises intervals within the cluster and the last sub-period (post/ending critical) comprises post-cluster intervals. In patients 1, 6 and 8, in which the critical phase was attached to the seizure onset, the last interval was considered to belong to the post/ending critical. Error bars denote SEM (standard error of the mean). Stars denote that there was a significant difference between the critical and the post/ending-critical sub-periods of the pre-seizure period (P<0.05, Wilcoxon test)

As observed earlier, the critical phase was not in general attached to the seizure onset of every patient. Hence, how could the critical phase be related to earlier reported evidences on the pre-ictal state? To address this question, we divided both recording sessions into the critical phase, and sub-periods immediately before (pre-critical phase) and after (post/ending critical phase) the critical phase (Fig 3C, Fig S3C for control patients). For those patients with critical phases attached to the seizure onset (Patients 1, 6 and 8) we considered the post-critical phase to comprise the last window time samples of the critical phase. In each sub-period we evaluated the mean functional connectivity during both recording sessions. Fig 3C shows that the mean connectivity exhibited a non-significant increase during the critical phase of the pre-seizure period (Fig 3C, P>0.2, paired Wilcoxon test, n=7 patients). In contrast, when comparing the critical and the post/ending-critical phases of the pre-seizure period, the mean connectivity decreased significantly over all patients (Fig 3C, P=0.02, n=8 patients) in concordance with previous works (Mormann *et al*. 2003, Le Van Quyen *et al*. 2005, Stacey *et al*., 2011). This result was validated at a single-patient level in 7 out of 8 main patients (Fig S6). Importantly, the post-critical effect was not present during the control period (P>0.9), suggesting that the global connectivity decrease was specific of the preseizure period and could be driven by the critical phase.

### Reduced network state variability spans across spatial and temporal domains

As introduced earlier, the centrality entropy quantified the (spatio-temporal) variability of simultaneous centrality sequences in a single scalar value. Then, how was the variability reduction individually expressed along recording sites and along time samples? To answer this question, we repeated the previous non-parametric statistical analysis (Fig 3A) over both recording periods using the spatial and temporal versions of centrality entropy independently (*Materials and Methods*). Fig S7B shows that the statistical effect was present in both dimensions for every patient but it was not equally distributed over space and time in all cases. In sum, the decrease of network state variability observed during the pre-seizure period was associated with the occurrence of more similar centrality values over time (less temporal variability), which in general exhibited more homogeneous centrality values across recording sites (less spatial variability).

### Altered occurrence of high-connectivity states explains reduction of variability

The previous results described that network states (as modelled by the eigenvector centrality measure) became more temporally redundant and more spatially homogeneous during the critical phase. In turn, this reduced variability was associated to a non-significant variation of the mean connectivity across patients (Fig 3C). Yet, how was the actual interplay between network dynamics and connectivity alterations during the pre-seizure period? An initial time-varying analysis of the mean functional connectivity (averaged over all recording sites’ pairs) did not reveal consistent and sustained cross-period differences over patients (Fig S9). We then related the reduction in network variability to alterations in the occurrence of certain states. In particular, were there specific time-varying states producing the reported effect? We here explored this question and inspected the eigenvector centrality sequences during the control and pre-seizure periods. A visual inspection on these vector sequences for every patient suggested the hypothesis that the amount of “homogeneous states” (represented as yellow strips in the plot) was larger during the pre-seizure period than in the control period. Interestingly, these homogeneous states were specifically associated with high-connectivity correlation matrices in most of the patients (Fig 4A).

**Fig 4:**
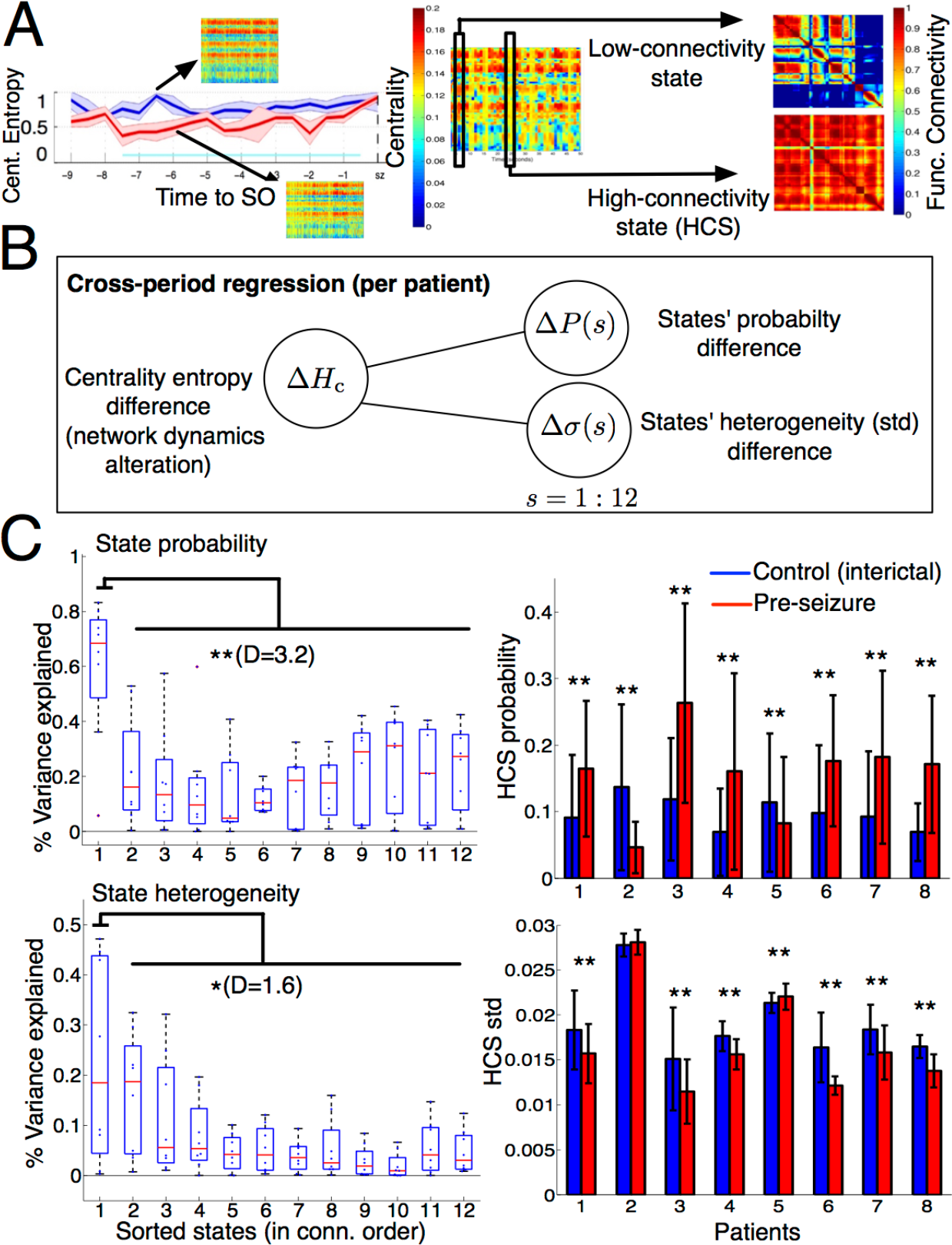
High-connectivity instances influence network dynamics alterations. (A) Inspection of centrality values around the critical phase (in cyan) suggested a higher presence of homogeneous (yellow strips) values across recording sites during the pre-seizure period (left), which were associated to high-connectivity matrices (HCS, right). Color intensity (blue=lowest, red=highest) represents centrality and connectivity values across recording sites. (B) Schematic representation (one per patient) of crossperiod entropy differences as a function of two families of regressors: changes of (discretized) state probabilities and changes of state homogeneities across recording sites. (C) Variance explained by each family of regressors (Top, state probabilities; bottom, state homogeneities) in every patient highlights high-connectivity states as a common putative driver of the critical phase. Left: For each patient, discretized states (n=12) were sorted along the horizontal axis in mean connectivity decreasing order. For each sorted state, boxplots show the distribution of the coefficient of determination (%variance explained) of each state across patients. Stars (* P<0.05, ** P<0.01, Wilcoxon test) denote the significance and D (Cohen’s *d*) denotes the effect size of the difference between the coefficients of determination of HCS and the remaining states. Right: Cross-period comparison of regressors values associated with high-connectivity state (HCS) during the critical phase between the control and the pre-seizure period. Bars denote the average value of each regressor during the critical phase of the preseizure (red) and control periods (blue) per patient. Error bars denote one standard deviation. Upper stars show that the differences in HCS probabilities and homogeneity were significant in all studied patients (* P<0.05, ** P<0.01, paired t-test) after multiple-test correction. All variables in this regression analysis were computed in time windows of 200 time samples (120s).

Centrality vector sequences like the one presented in Fig 4A were observed to be recurrent over time. Then, we used a clustering algorithm to extract the 12 most representative vectors over both periods of interest and classified each centrality vector at any given time accordingly (*Materials and Methods*). Consequently, the sequence of centrality vectors turned into a sequence of discrete states whose frequency over any time interval (probability) could be computed and compared across control and pre-seizure periods. Then, we formally tested the hypothesis that the larger presence of homogeneous states during the pre-seizure period was associated to the observed reduction in network state variability in each patient. For each patient, we linearly regressed the cross-period centrality entropy difference over two independent state regressors: state probability and state heterogeneity (measured via the standard deviation across recording sites of the same state) differences (Fig 4B). We then computed the variance explained by each regressor via its coefficient of determination (R squared). To investigate the group-level influence of every state’s connectivity into these associations, states were sorted for each patient in decreasing order of connectivity (i.e., mean connectivity of its associated correlation matrix), and coefficients of determination linked to state probability (Fig 4C, top) and state heterogeneity (Fig 4C, bottom) differences were distributed in boxplots for each state. Interestingly, Fig 4C (left) shows for both regressors (state probability and state heterogeneity) that the most influential states on the reduced variability effect were those with largest connectivity associated matrices Specifically, the difference between the variance explained by the highest-connectivity states and the remaining ones was significant in both state probability (P<0.001, Wilcoxon test) and state heterogeneity (P<0.05) with large effect sizes (D=3.2, D=1.6, Cohen’s d). Then, we computed the Spearman correlation between the highest-connectivity regressors and the centrality entropy reduction to unravel group-level correlation trends. Correlation values were of r=0.63 (P<1e-5) and r=-0.45 (P<1e-5) for state probability and heterogeneity increases respectively indicating that the reduction of network variability was mostly explained by an increase in the frequency rate and homogeneity of the highest connectivity states.

To further investigate the interplay of high-connectivity states with the pre-seizure period, we evaluated cross-period state probability and heterogeneity differences at a patient level during the critical phase previously identified in Fig 3A (Fig 4C right). First, we found that the probability of HCS was significantly different in all patients across both periods (paired t-test, P<0.01, multiple test corrected, D>0.5). In 6 out of 8 patients HCS occurred significantly more often during the critical phase while they were less frequent in the remaining patients (Patients 2 and 5). Second, the homogeneity of HCS was significantly increased in most of the patients (paired t-test, P<0.01, multiple test corrected, D>0.5), except in Patient 5 were it significantly decreased, and in Patient 2 where it remained statistically equal (P>0.05). Although the influence of HCS into the pre-seizure period was consistent across all patients, the differentiated trends found in some specific patients (Patients 2 and 5) suggest that this influence might be modulated by context-dependent variables. In sum, HCS strongly contributed to make state dynamics less variable over time by reducing the occurrence of alternate states and imposing homogeneous centrality values across recording sites.

The key influence of HCS into pre-seizure dynamics prompted us to evaluate the underlying traces of iEEG data during their corresponding time instances in periods of high and low centrality entropy. Our inspection of iEEG data from distinct epileptogenic sites over sequences of HCS and nHCS instances (See Fig S11 for an example) identified these states as time segments where the electrical fields became transiently (low centrality entropy epoch) or more persistently (high centrality entropy epoch) synchronized. This synchronization was manifested through diverse patterns of oscillatory activity, which often included a slow wave. In parallel, a clinical evaluation by the epileptologists discarded any stereotyped epileptiform activity.

### Cross-validation analysis in additional interictal periods

We identified network dynamics changes in the pre-seizure period that were consistently expressed with a similar trend (sustained variability reduction) across a heterogeneous cohort of patients (Fig 1). Critically, these time-dependent changes could be associated to a common factor in all patients, namely, an alteration of recurrent high-connectivity time instances (0.6s) across recording sites. However, was this characterization specific of the pre-seizure period? Or could be alternatively ascribed to a post-implantation effect? To shed light into these questions, we analyzed additional 121 hours of interictal activity in 6 patients from time-matched periods that were placed two days before the seizure (‘pre-control’ period) and a varying number (across patients, mean=3,83) of days after the seizure (‘post-control’ period). These new interictal data was introduced in the analysis as schematized in Fig 5A. As control experiments, we defined two additional time-matched comparisons: a comparison between the pre-control and control periods (‘C1’) and a comparison between the seizure and post-seizure period (‘C2’). These new comparisons were then confronted with the original comparison particularized to the 6 patients. The overall analysis was made under the condition that period lengths were time-matched and balanced across comparisons for each patient. First, for every comparison, we repeated the non-parametric statistical analysis of Fig 3A to determine the existence of putative critical phases in other periods. While comparison C2 only yielded one patient with a significant effect, C1 revealed that entropy reductions could also occur in non pre-seizure periods in 5 out of 6 patients (Fig 5B). Nonetheless, when grouping the six patients in Fig 5C, the sub-periods found with C1 were not followed by a post-critical functional connectivity decrease, which was present in C0 as a significant trend (P<0.1, Wilcoxon test). Finally, we repeated the regression analysis of Fig 4B in patients with significant entropy reductions of C1 (five patients) and C0 (original comparison in six patients) and represented the results along analogous lines for each comparison. Crucially, for C1 periods, the variability decrease was more weakly explained by cross-period HCS differences than in C0 periods. Indeed, the significant trend in the gap between the variance explained by HCS states and the variance explained by non-HCS states in state probability (P<0.1, D=1.5) and in state homogeneity (P<0.1, D=1.6) in C0 could not be reproduced in C1 (P>0.1). Although decreases in network state variability may occur across consecutive days (C1) preceding a seizure, we provided evidence that those occurring during the pre-seizure period were specifically tied to high-connectivity states alterations and a subsequent functional connectivity decrease.

**Fig 5:**
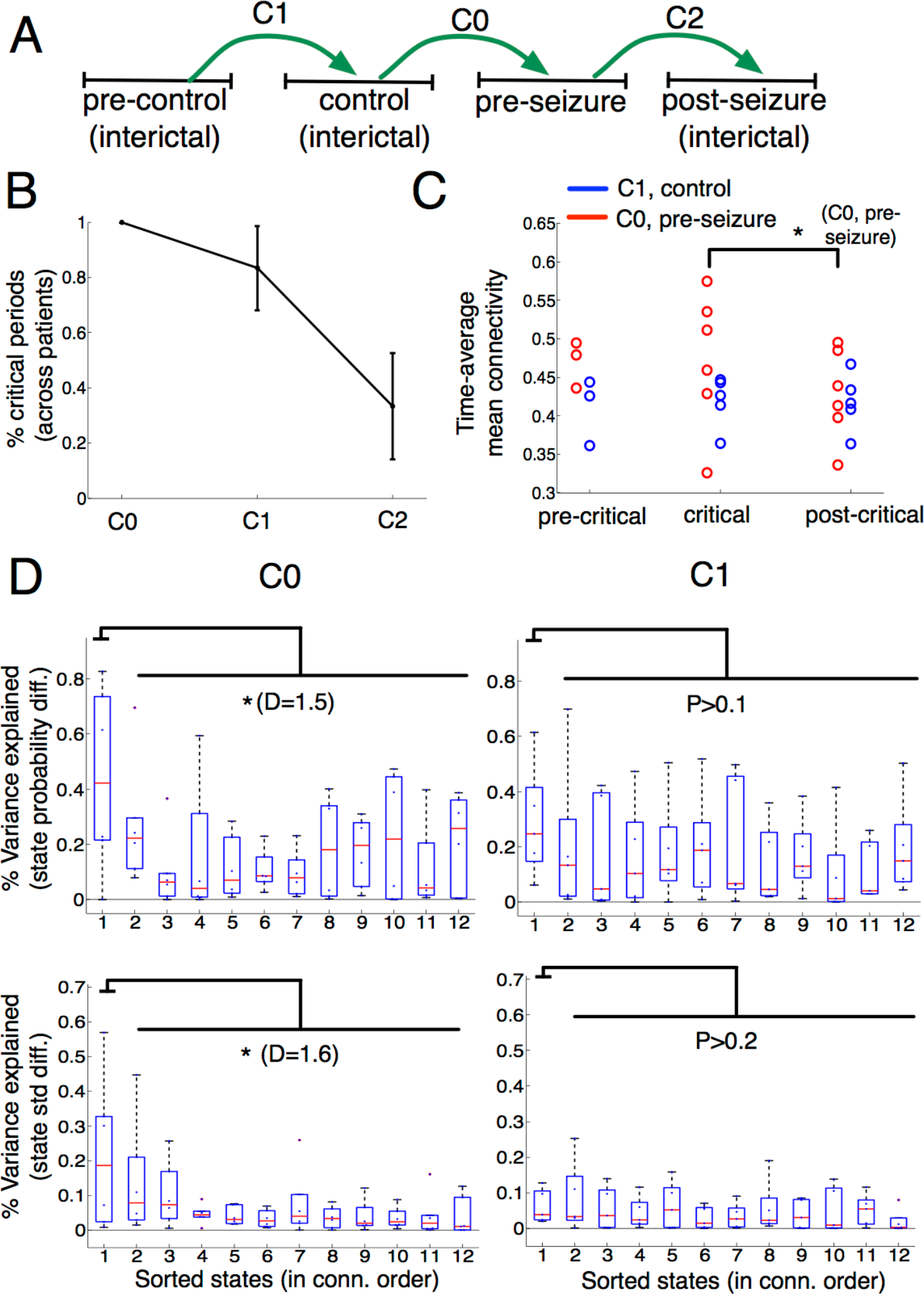
Cross-validation analysis in additional periods (Patients 2-6, and 8) (A) Schematic representation of the cross-validation analysis involving Patients 2-6, and 8, and periods of the same lengths (within patient) for the 4 periods. Time-matched periods from 2 days before the seizure (pre-control) and from a varying number of days after the seizure (post-seizure) gave rise to two additional cross-period comparisons (C1 and C2) to the previously analyzed (C0). (B) Percentage of significant intervals across patients in the cross-period comparisons C0-2 (cluster-based test, P<0.01). Errors bars indicate SEM. (C) Reproducing the same pre-seizure connectivity curve in Fig 3C via comparison C0 (in red) and C1 (in blue). The upper star indicates that the difference between the time-average connectivity values in the critical and post-critical phase trended a significant effect (* P<0.1, Wilcoxon test, N=6). (D) Variance explained by each family of regressors of Fig 3C using the comparison C0 (left) and C1 (right). The upper star indicates that the difference between the coefficients of determination of HCS and the remaining states trended a significant effect (* P<0.1, Wilcoxon test). D denotes the effect size (Cohen’s *d*) of this difference.

### Influence of the critical phase into epileptogenic sites

Importantly, network dynamic changes observed during the pre-seizure period could be associated to an altered occurrence of HCS in all patients. Yet, how this seemingly physiologic alteration could evolve into generating seizures? In particular, how was this effect manifested in those regions that were involved in seizure generation? To further relate our findings to the ictogenesis process, we particularized our analysis to the clinically mapped epileptogenic sites of two patients with very good post-surgical outcome (Engel I) and a follow-up period of more than four years (Patients 1 and 3, Fig 1, *Materials and Methods*). Both patients are seizure free (Engel 1) with Patient 3 exhibiting some residual ictal symptomatology (seizure auras). In these patients, we specifically investigated the influence of epileptogenic sites in the pre-seizure network dynamic changes. To provide a complete comparison of sites, we independently analyzed seizure-onset zone sites (SOZ, brain zone involved in the initial stages of the seizure spread), resected zone sites (RZ, brain zone that rendered seizure-freeness after its resection) and the remaining sites (nEZ, nonepileptogenic sites). In both patients, we note that SOZ was not fully included in the RZ and thus, SOZ and RZ were partially overlapping regions. To carry out this region-specific analysis, we first evaluated the temporal mean and standard deviation of the recording sites’ centrality in the SOZ, RZ, and nEZ sites over the control and pre-seizure periods. Fig 6A plots the time-average centrality of RZ and nEZ as a function of the remaining time to seizure onset. Interestingly, this figure illustrates in both patients that the time-average centrality of the RZ was higher than the nEZ over each period of interest and during the critical phase (in cyan) the centrality of RZ sites was reduced at the expense of an increase in the centrality of nEZ sites. This preliminary observation suggested that both regions could participate in the pre-ictal dynamics. However, was this participation equal across the three considered regions? Fig 6B characterizes the network dynamics of the three regions by comparing the temporal standard deviation of their recording site’s centrality in SOZ (inner left), RZ (inner central) and nEZ (inner right) regions for control (blue) and pre-seizure (red) period, inside (outer left) and outside (outer right) the critical phase. To assess cross-period differences across regions of variable size we highlighted significant differences (P<0.05, paired t-test, multiple-test corrected) exceeding an effect size threshold of 0.5 (large effect, Cohen’s d). Using this quantification, Fig 6B shows that the largest decrease in the centrality variability (D>0.5) of Patient 1 was only localized in the RZ during the critical phase. For Patient 3, large effect sizes were found in RZ but also in nEZ during the critical phase. Outside the critical phase, cross-period differences attained lower effect sizes (D<0.3).

**Fig 6:**
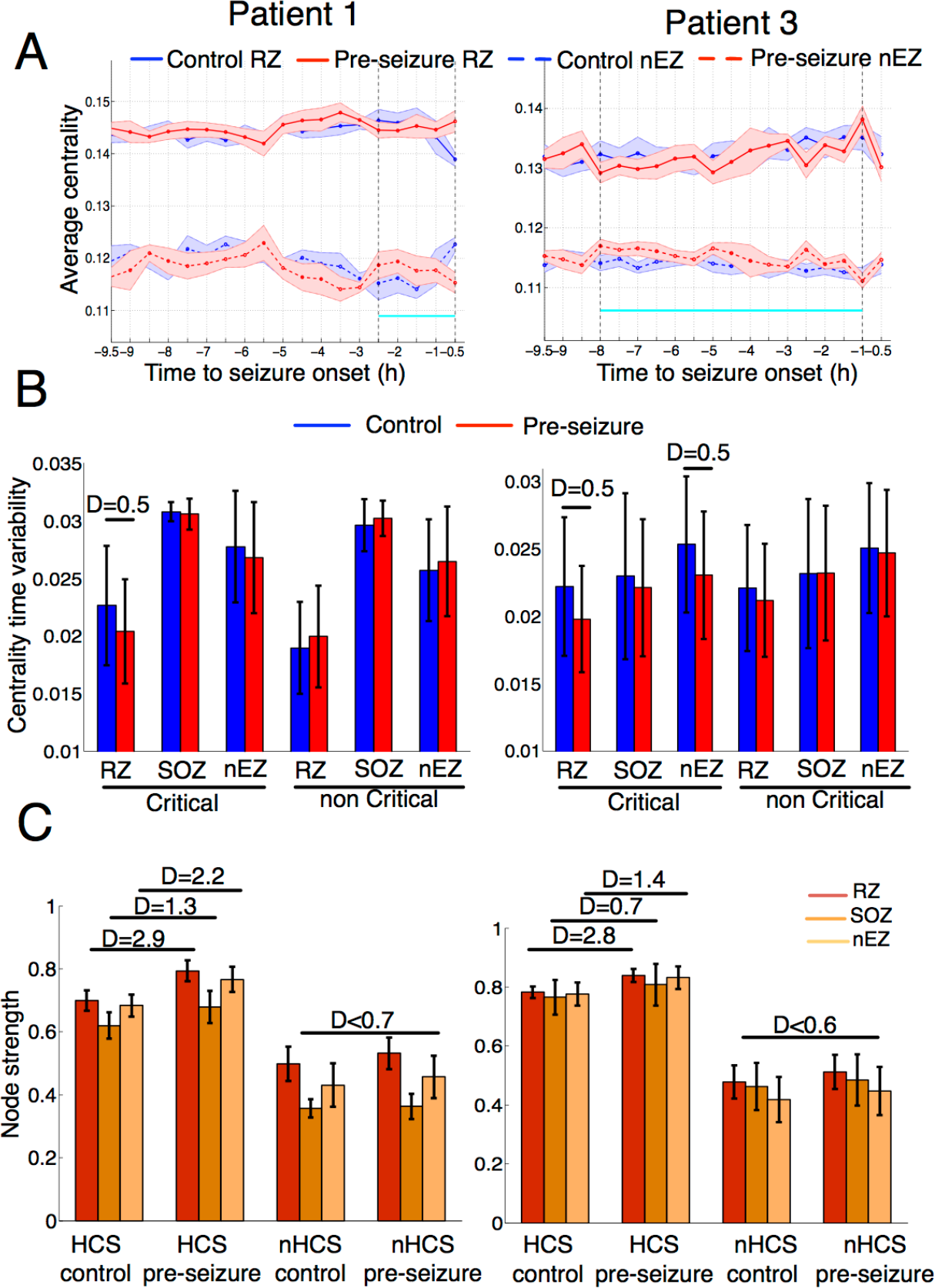
Epileptogenic sites are specifically altered in the critical phase (Patients 1 and 3). In the two patients with best post-surgical outcome after resectomy, recording sites in the resected zone (RZ), seizure onset zone (SOZ) and in none of these regions (nEZ) were independently analyzed (*Materials and Methods*). (A) For each patient and period, site-average eigenvector centrality in the RZ, and nEZ averaged within non-overlapping and consecutive time windows of 120s (200 samples) during 9.5 hours prior to seizure onset time. In solid line, average centrality of the RZ. In dashed line, average centrality of the nEZ. Blue and red curves stand for the control and pre-seizure periods respectively. For illustration purposes, curves were averaged within windows of 30 minutes (15 samples per window) to enable direct comparison with the estimated critical phase (highlighted in cyan between two dashed vertical lines). Error bars denote one standard deviation. (B) Cross-period comparison (control in blue, pre-seizure in red) of sites’ centrality variability averaged over RZ, SOZ and nEZ inside (critical, left) and outside (non critical, right) the estimated critical phase (cyan segment in A). Each sample per recording site was computed by performing an average (across pre-ictal and non critical phases) of the centrality’s temporal standard deviation measured in non-overlapping and consecutive time windows of 120s (200 samples). (C) Effect of high-connectivity state into the epileptogenic zone. For each patient, bars showing the site-average connectivity strength RZ, SOZ and nEZ during the high-connectivity clusterized states (HCS, outer left) of both patients and during the remaining states (nHCS, outer right) in control (inner left) and pre-seizure (outer left) periods within the critical phase. Strength samples were computed for each site by performing averages over each set of time instances (HCS and non HCS) during the critical phase. In (B) and (C), sizes of significant effects (paired t-test, P<0.05, multiple-test corrected) equal or larger to 0.5 were reported using Cohen’s *d* and approximated to the first decimal. In all subfigures, error bars represent ± one standard deviation.

We next investigated the influence of HCS on epileptogenic and non-epileptogenic sites to further describe the functional alterations occurring during the critical phase. More specifically, we compared the average connectivity per site (node strength) in the RZ, SOZ and nEZ during the presence of the HCS and the remaining states (nHCS) in each patient for control and pre-seizure periods in the critical phase (Fig 6C). This analysis revealed several findings. First, in both patients cross-period differences in strength occurred more prominently during HCS (average D >1.8) than in nHCS (average D <0.7). Second, during HCS, strengths increased from control to pre-seizure periods consistently in the three studied regions while the differences were of varying sign across regions during nHCS. Third, the region that exhibited the highest increase in strength was the resected zone for both patients (D=2.9, 2.8), followed by the non-epileptogenic sites (D=2.2, 1.4) and the seizure-onset zone (D=1.2, 0.6). Hence, the abnormal occurrence of HCS altered the connectivity gradient between epileptogenic and non-epileptogenic regions by strongly boosting the connectivity of the RZ sites. In particular, during the critical phase of the pre-seizure period, this increased connectivity was more persistent than in the control period resulting in a reduced variability of RZ centrality values (Fig 6B). We finally evaluated in both patients how the post-critical functional connectivity decrease (Fig 3C) was spread over the three regions. Fig S12 shows that this effect was reproduced in each region (D≥0.9) with epileptogenic sites showing a more prominent decay (average D=1.7) than non-epileptogenic sites (average D=1.35).

To relate some our regional findings with the patients’ post-operative outcome, we extended the analysis of the sites’ temporal variability (Fig 6B) to the main patients’ entire cohort (Fig S13). This included three patients that underwent RFTH with variable outcomes (Patients 2,6, Engel I and Patient 8, Engel III), two patients with bad post-surgical outcome after a follow-up period of more than one year (Patients 4 and 5, Engel III) and one patient that was seizure-free after SEEG monitoring (Patient 7). The results are depicted for each patient in Fig S13 and summarized in Fig S14, providing preliminary evidence that bad post-operative outcomes could be associated with non-resected and non-ablated sites exhibiting pre-seizure alterations (Fig S14).

## Discussion

This study examined the existence of a common alteration principle in brain network dynamics during long-lasting periods of activity preceding the first clinical seizure in 10 patients with focal refractory epilepsy. Using a comparative analysis between genuine preseizure periods and time-matched periods of interictal activity per patient, we were able to consistently show a sustained decrease in the variability of network states that was followed in most of the patients by a functional connectivity drop of approximately 30 minutes before the seizure onset (Fig 7). Further analysis revealed factors altering this variability in the temporal (time samples) and spatial (recording sites) domains. First, this decrease in network variability was associated with an abnormal occurrence of high-connectivity states during pre-seizure periods as compared to previous days. Second, the reduction in temporal variability was mainly localized in the resected zone of two patients with best post-surgical outcome.

**Fig 7:**
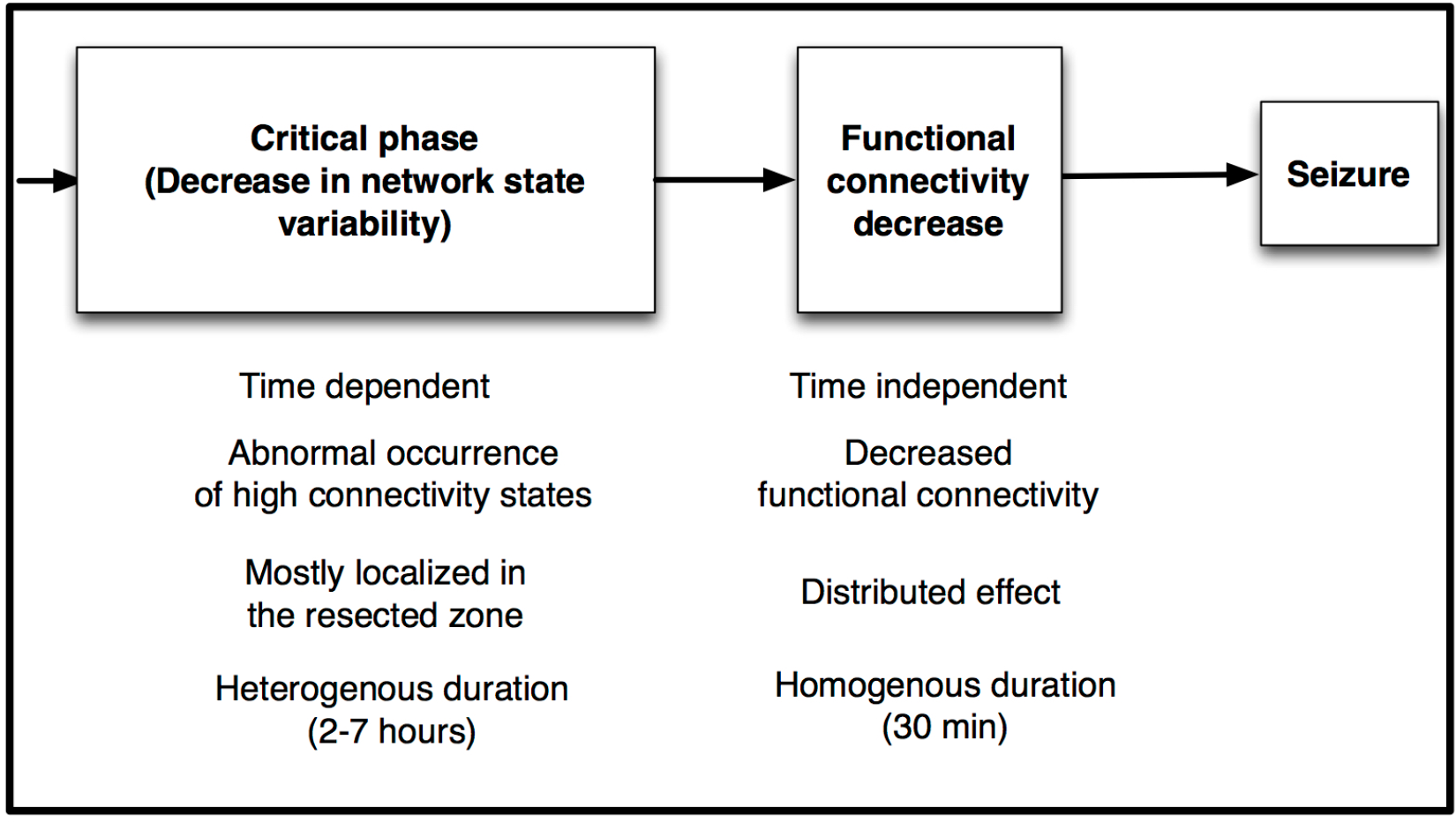
Scheme representing the pre-ictal characterization with two sequential events of different nature and duration: the critical phase and the global functional connectivity decrease.

Over the last decade, fMRI studies have showed growing evidence that dynamic connectivity patterns (“brain dynamic repertoire”) may be an intrinsic property of brain function and disease (Hutchinson *et al*., 2013). Particular examples of disrupted dynamics have been found in Alzheimer’s disease (Jones *et al*., 2012) and neuropsychiatric disorders (Damaraju *et al*., 2014) whose translation to new clinical biomarkers is still matter of discussion (Deco and Krigelback 2014, and references therein). In modern epilepsy research, the dynamic principle of brain function has been postulated to be commonplace to understand the ictogenesis process (Richardson 2012) but most network studies have studied alterations in static functional network paramters with a few recent exceptions (Kuhnert *et al*., 2010, Dimitriadis *et al*, 2012, Kramer *et al*., 2011, Burns *et al*., 2014, Morgan *et al*., 2015, Khambhati *et al*., 2017). In this context, our approach differs from previous works in several key elements. To name a few, the formulation of an hypothesis about the variability of functional network states at short time scales (rather than using a grand-average measure), the analysis of long-lasting (∼10h) continuous interictal periods (rather than a selection of short epochs), and more importantly, the use of time-matched reference epochs outside the pre-seizure period to assess specificity. We next elaborate further on the latter point.

When studying the variability of brain dynamics along long recording periods, one is confronted with the confounding effect of circadian rhythms (Kuhnert *et al*., 2010, Rocamora *et al*., 2013, Geier *et al*., 2015a), which span across sleep and wake phases. These rhythms may become critical when one characterizes specific brain configurations associated with the pre-ictal state, which has been shown to approximately last 4 hours (Mormann *et al*., 2007). Previous studies on the pre-ictal state have analyzed pre-ictal changes with reference to previous interictal periods, not necessarily time-matched. Inspired by a previous work (Andrzejak *et al*., 2003), the strategy used here tackled this issue by defining time-matched reference periods from precedent and subsequent days, thus allowing for a more specific identification of pre-ictal changes in brain network dynamics. Although this approach may not be sufficient to control for all daily physiological state transitions, our preliminary results on the relationship between patients’ putative critical phases and seizure onset times discard the influence of daily rhythms into our main results. However, a larger cohort of patients with variable seizure times and a good readout of their sleep phases will be necessary to address this question in the future. Another key aspect of the study was the use of the first monitored clinical seizure occurring during the first implantation days. This choice was pivotal to analyze comparable long-term network dynamic changes across patients with limited influence of confounding factors such as the reduction of antiepileptic drugs, the effect of previous ictal processes and the response to clinical stimulation. In most of the studied patients, this first seizure was the first event of a succession of seizures separated by short interictal periods of a few hours or minutes, which are clinically known as seizure clusters (Rose *et al*., 2003). Hence, understanding the pre-ictal process of this initial seizure can also have important consequences for the control of later correlated events. In any event, the analysis introduced here should be extended to subsequent seizures in future studies to determine whether the presented characterization is specific of seizures preceded by long interictal periods.

A central question in seizure prediction research has been the role of synchronization (Jiruska *et al*., 2013) during the pre-ictal period. Some studies have reported drops in synchronization a few hours before seizure onset (Mormann *et al*., 2003) while others have pinpointed the coexistence of distinct synchronization states depending on the recorded structures (Le Van Quyen *et al*., 2005, Van Mierlo *et al*., 2014). Even though a clear mechanism of such alterations is still missing, the most successful algorithms applied to large data sets make use of correlation matrices as key data features (Binkmann *et al*., 2016). The findings presented in this study support the view that pre-ictal correlation patterns are state dependent (Le Van Qyuen *et al*., 2005, Takahashi *et al*., 2012, Khambati *et al*., 2017) over time windows of 600ms and hence, their alterations should be analyzed and interpreted at this time scale. More precisely, our results suggest that a time-dependent variation in the occurrence of highly correlated time instances may be at the origin of the pre-ictal state. This variation was manifested in most of the patient as an excess of high-connectivity states, while in two patients it was manifested as a deficit. Although pre-ictal connectivity trends are known to be patient-specific (Jiruska *et al*., 2013), they should be further investigated against the influence of patient-dependent variables (e.g., implantation schemes, monitored behavioural states), a question that was out of the scope in this study.

In recent years, there has been accumulated evidence that seizure generation and spread involves complex interactions between seizure-generating and surrounding areas (Rummel *et al*. 2013, Khambati *et al*., 2015, Khambati *et al*., 2016). Evaluating network dynamics in patients with good post-surgical outcome (>4 years), we were able to relate our findings to clinically mapped epileptogenic sites, namely the seizure onset zone and the resected zone, as well as the remaining sites. In these patients, the average contacts’ centrality was higher in the epileptogenic sites for the entire analyzed periods in line with previous studies (Wilke *et al*., 2011, Van Mierlo *et al*., 2013). However, this gap was mainly produced by sites in the resected zone that were not part (but in nearby regions) of the seizure onset zone (Fig 6A and 6C), also in concordance with a recent study (Geier *et al*., 2015b). Not surprisingly, changes in this average centrality level within periods occurred during the critical phase where centrality values from both regions approached (Fig 6A). Crucially, this centrality change was accompanied by a significant decrease in the centrality (temporal) variability of the resected zone (Fig 6B), which was specific in Patient 1 and also present in the nonepileptogenic sites in Patient 3, who presented a slightly worse post-surgical outcome. The analysis on the influence of high-connectivity states into validated epileptogenic sites provided initial evidence that these states might destabilize physiological state dynamics by increasing the connectivity of key sites within the epileptic network (resected zone) during the critical phase (Fig 6C). The consequence of this phase is shown to be a global functional connectivity decrease, which is more prominently manifested across epileptogenic nodes (Fig S10). We speculate that this decrease in connectivity could be the result of critical parts of the epileptic network adopting a more autonomous activity that would result in the generation of a seizure. The extension of our analysis to additional patients with different post-operative outcome suggest that pre-seizure alterations in centrality variability may be a promising biomarker of targetable epileptogenic regions during surgery and ablation (Fig S14). Yet, a larger study including more seizure-free patients will be necessary to fully elucidate the mutual influence of physiological network dynamics and the epileptic network during the transition from interictal activity to focal seizures.

The results shown in this study prompt to introduce new ingredients in seizure-prediction algorithms such as the control for daily rhythms (Karoly *et al*., 2017) and the continuous tracking of time-dependent linear connectivity alterations at short time scales (<1s). Some considerations are yet to be mentioned. First, the use of intracranial recordings is a limiting factor in the spatial analysis of brain states, thus making them a priori subject-dependent. Nonetheless, it is recognized that the SEEG methodology offers an optimal temporal and spatial resolution of neurophysiological recordings for neural signal analysis in comparison with other techniques in patients with epilepsy. Second, this study was aimed at defining network states in a linear and instantaneous form using zero-lag functional connectivity rather than effective connectivity (Friston 2011). Although our results were validated against a non-linear coupling measure at different narrow bands, the extension of our analysis to non-linear (Tauste Campo *et al*., 2015) and linear (Gilson *et al*., 2017) directional methods in follow-up studies may provide additional information on specific connectivity changes underlying preseizure alterations. In conclusion, this work provides electrophysiological evidence for characterizing the pre-seizure period as a long-lasting process in which epileptic networks undergo a sequential functional reorganization. Further investigations under this conception will help unravel seizure generation mechanisms from a network perspective, provide practical insights into how to predict and control ictal activity, and may constitute a general approach to analyze dynamic alterations of other neuropathologies.

## Materials and Methods

### Patients and recordings

A total number of 344 hours of SEEG recordings from ten patients with pharmacoresistant focal-onset seizures were analyzed. A summary of the patients’ characteristics is given in Fig 1. We included patients who presented the first spontaneous clinical seizure in a time frame that allowed us to perform a controlled analysis of EEG recordings during the pre-seizure period. Specifically, each patient in the study was selected if her first video-SEEG monitored clinical seizure had occurred after at least 30 hours (average value: 71.4±19.1 hours; mean±std) with no presence of spontaneous clinical seizures. Among the selected patients we included two patients presenting potential perturbation factors affecting the pre-seizure period (Patients 9 and 10). Patient 9 had been electrically stimulated 16.5 hours before the first recorded seizure and Patient 10 presented a subclinical seizure 6.1 hours before the first clinical seizure onset.

For each patient, the selection of recording sessions was as follows. We considered up to 12 hours before the first monitored clinical seizure occurred. As a baseline reference, we selected the same time period from the previous day (control period). For independent validation of our results, we selected additional time-matched periods of variable length in 6 patients (Patients 2-6 and 8, average period length: 10 hours) from two days before the seizure onset (pre-control period), and a few days after the seizure onset (post-control period, average value=3.83 days). No more patients could be added to the validation analysis for preictal time limitations (Patients 7 and 10), a substantial modification on the implantation montage during the first monitoring days (Patient 1) or the presence of direct electrical stimulation sessions in the iEEG (Patient 9).

After detecting recording cuts in a few patients, we restricted the analysis to 11 hours per session in patients 1-9 and to 2.4 hours per recording session in Patient 10 to ensure a time-matched cross-period comparison. Among the selected patients, two patients achieved seizure freedom after surgical resection and radiofrequency thermocoagulation (RFTC, Cossu *et al*. 2015) with a follow-up of 3 years and 2 years respectively (Patients 1 and 2, Engel 1A). An additional patient only exhibited seizure auras after surgical resection and a follow-up of 3 years (Patient 3, Engel 1B). We considered Patients 1 and 3 to have a validated very good post-surgical outcome. Hence, for the purpose of analyzing epileptogenic sites, we separately considered the diagnosed seizure onset zone and the resected zone of these two patients. The seizure-onset zone was independently marked by two epileptologists (AP and RR) and consisted of n=5 (anterior hippocampus) and n=9 (anterior hippocampus, amygdala) recording sites for Patient 1 and 3 respectively. The resected zone covered 24 contacts in Patient 1 (parts of anterior hippocampus, temporal pole and entorhinal cortex) and 12 contacts in Patient 3 (parts of anterior, posterior hippocampus, and amygdala). The remaining patients presented one of these cases: they had not undergone surgery (Patients 2, 6, 8, 9), had a non-sufficiently long follow-up period (<6 months, Patients 4 and 5), had not been yet operated (Patient 7) or exhibited a bad post-operative outcome (Patients 10). All recordings were performed using a standard clinical EEG system (XLTEK, subsidiary of Natus Medical) with a 500 Hz sampling rate. A uni- or bilateral implantation was performed accordingly, using 5 to 15 intracerebral electrodes (Dixi Médical, Besançon, France; diameter: 0.8 mm; 5 to 15 contacts, 2 mm long, 1.5 mm apart) that were stereotactically inserted using robotic guidance (ROSA, Medtech Surgical, Inc).

### Data pre-processing

EEG signals were processed in the referential recording configuration (i.e., each signal was referred to a common reference). The sets of electrodes included in this analysis are reported in Fig 1 and displayed in Fig 1 (top row). All recordings were filtered to remove the effect of the alternate current (Notch at 50 Hz and harmonics using a FIR filter). Then signals were further band-pass filtered between 1Hz and 150 Hz to remove slow drifts and aliasing effects respectively. Artifacts were removed in each period by detecting time window samples (600ms) where mean (over pairs of sites) correlation values and mean (over sites) voltage amplitudes were 3 standard deviations larger than the median values across each period. To perform functional connectivity analysis each EEG signal was divided into consecutive and non-overlapping 0.6s-long windows (300 samples with 500Hz sampling rate) to balance the requirements of approximate stationarity of the time series (requiring short epochs) and of sufficient data to allow accurate correlation estimates (requiring long epochs).

### Functional connectivity analysis

There are different methods to assess functional connectivity from time series data based on coupling measures (Pereda *et al*., 2005, Wendling *et al*., 2009). Previous research on the comparison of linear and non-linear coupling measures has resulted in having distinct “ideal” measures for distinct studied situations (Stefan *et al*., 2013). Here we chose to employ a zero-lagged linear correlation measure for its good tradeoff between simplicity and robustness (Wendling *et al*., 2009) and more importantly, because it allowed for a convenient definition of *network state* as it will be explained later.

Leat X and *y* be two *N*-length time series representing two recorded signals and let 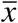 and 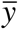 be their respective sample means. Their sample (Pearson) is estimated as

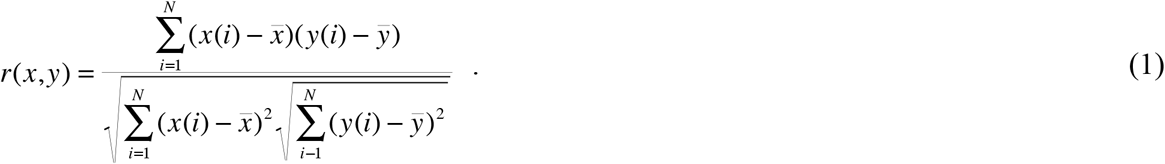

For each patient and each consecutive 0.6s-long window we computed the absolute value of the coupling measure across all pairs of electrode contacts. For most of the patients, the overall pairwise computations resulted in approximately 123000 sequential connectivity matrices combining both recording sessions (control and pre-seizure periods). In the current study, we did not test the statistical significance of each pairwise coupling since our purpose was to track the overall network dynamics regardless of pairwise thresholding methods.

### Definition of network states

For each patient, we characterized each correlation matrix as a functional network. This network was modelled as a weighted undirected graph, where electrode contacts represented the nodes and pairwise correlation values across represented their weighted edges (Ponten *et al*., 2007). Then, we computed the network measure of eigenvector centrality for each connectivity matrix (Newman, 2010). For a given graph *G=(V,E)*, let *A=(a_v,t_)* be its weighted adjacency matrix. The relative centrality score *x_v_* of vertex *v* can be defined as

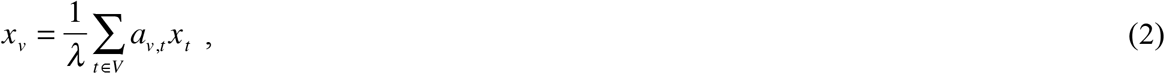

which can be rearranged in a matrix form as *λx = Ax*.

Given the requirement that all entries in *x* must be non-negative, the Perron-Frobenius theorem implies that only the greatest eigenvalue results in a proper centrality measure (Newman 2010). Hence, the centrality measure is given by the eigenvector associated with the largest eigenvalue of the connectivity matrix. Then, the *i*th contact is assigned the *i*th component of this eigenvector where *i* goes from 1 to number of recording sites in a patient. The eigenvector centrality is by definition a self-referential measure of centrality, i.e., nodes have high eigenvector centrality if they connect to other nodes that have high eigenvector centrality (Rubinov and Sporns, 2010), which ultimately provides a measure of relative importance of each node in the network. The eigenvector centrality measure has been applied to resting-state fMRI studies (Lohmann *et al*., 2010) and more recently to ECoG recordings of epileptic patients (Burns *et al*., 2014).

By computing the centrality in each 0.6s-long connectivity matrix we obtained for each patient independent eigenvector centrality sequences along each recording session. If we consider each connectivity matrix to represent a brain state (Allen *et al*., 2014), these vectors can be regarded as representative elements of these states in a vector space of dimension equal to the number of recording sites. Further, these vectors point to the direction that best summarizes the original brain state. In particular, every time that a significant change arises in the connectivity matrix, the eigenvector centrality rotates to update the relative importance (“centrality”) of each contact within the new network configuration.

### Choice of zero-lag correlation and eigenvector centrality

Computing the eigenvector centrality over zero-lag connectivity matrices was key to regard our network state measure as an informative summary of how the set of simultaneous iEEG recording were instantaneously coupled within a short time window. Indeed, under these conditions, the eigenvector centrality corresponds by definition to the first principal component of the (normalized) covariance matrix, i.e., the vector in the space of recording sites that accounts for the largest variance of the whole set of (normalized) iEEG recordings in a given time window. Combinations of other coupling measures and network features could lead to alternative definitions of network states. For the sake of comparison, we also provide in the Supplementary Information the results obtained by combining zero-lagged correlation with a different network feature, the node strength, which can be defined as the average pairwise connectivity of this node with the remaining ones (Rubinov and Sporns 2010, Khambhati *et al*., 2016). Fig S3 shows that the node strength yielded in general statistically weaker results than the eigenvector centrality. Further, we investigated the possibility of combining a synchronization measure such as the phase-locking value (Lachaux *et al*., 1999) with the eigenvector centrality. This measure may capture contributions of non-zero lag couplings as well as non-linear effects. To illustrate the difference between both measures in the frequency domain, we repeated the cluster-based statistical analysis of Fig 3A for consecutive frequency narrow bands of 4Hz (from 1 to 120). Fig S4 shows that the results were qualitatively similar across all bands for most of the patients. Yet, in those patients where discrepancies were found, the phase-locking value measure yielded weaker peaks than the zero-lag correlation.

### Evaluating network state dynamics via Gaussian entropy

Our goal was to evaluate the variability of these representative states in each period. The long sequence of centrality vectors for each period can be equivalently regarded as a stream of simultaneous centrality time series, one for each recorded contact. Then, one can evaluate the spatio-temporal variability of the centrality time series through the application of the multivariate Gaussian entropy (Cover and Thomas, 2012) in a given estimation time window that we choose for this study to be 120s. The multivariate Gaussian entropy is defined as

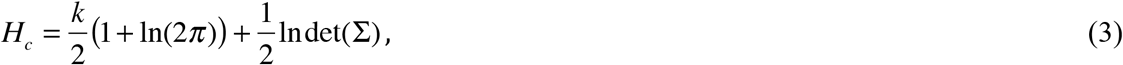

where k is the number of recording sites, and Σ is the covariance matrix of the centrality time series estimated in a the estimation windows. By considering centrality vectors to be independent, Σ in (4) becomes a diagonal matrix, and the Gaussian entropy captures the aggregated variability of the centrality vectors across the temporal dimension:

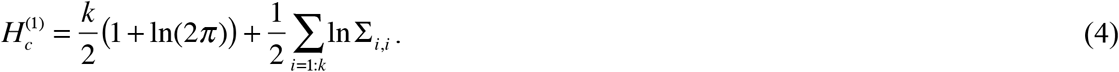

By subtracting (5) from (4), one can evaluate the variability of the centrality vectors across the spatial dimension:

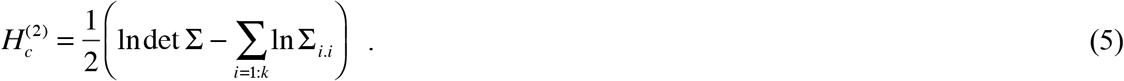

Hence, the two contributions sum up to give the Gaussian entropy (4):

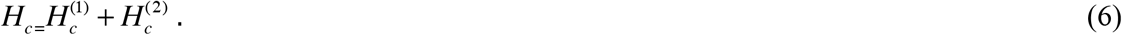

### Choice of window sizes (correlation and entropy)

The choice of 0.6s (300 samples) on the correlation window was critical to gain statistical power. Choices of 1, 5 or 10s were shown to weaken the detection of network dynamics changes because they were intermingling high and low connectivity effects in the same window. On the other hand, values of entropy windows ranging from 100 to 200s yielded quite stable results. We selected a window size of 120s (200 samples) because it offered a good tradeoff between estimation accuracy (200 samples are good enough to estimate covariance matrices of at most 120 variables) and stationarity.

### State clusterization

To associate the network variability decreased observed in all patients with the occurrence of specific recurrent connectivity states, we jointly clustered the eigenvector centrality sequences in the analyzed time-matched comparisons using the k-means algorithm (Forgy, 1965). We applied this clusterization in patients 1-9 where the number of eigenvector centrality samples was comparable. In the main results we fixed the number of clusters to 12 to cover a sufficiently wide range of visually inspected connectivity states per patient. This cluster size was selected after exploring the stability of the results illustrated in Fig 5 for the range of values n=8:12. In particular, Fig S9 shows that these results were qualitatively very similar for the choices n=8,10,12.

### Statistical analysis

The pre-seizure decrease in centrality entropy was statistically tested as follows. We started by windowing consecutive entropy samples (n=15, 30 minutes) in non-overlapping and paired time segments across each period and then we computed the effect size for each segment pair using Cohen’s *d* (Cohen 1992). We then clustered adjacent segments with a criterion of effect size being larger of 0.15 (low-medium effect) over a minimum of 4 adjacent segments (2 hours), and considered the aggregated sum of these segments’ effect sizes as the main statistic. We further checked the statistical significance of this value through non-parametric statistical testing based on Monte Carlo sampling (Maris and Oostenveld, 2007). More concretely, for each patient with time segments satisfying the above criterion, we computed 1,000 random permutations of the centrality entropy samples across both conditions (within pre-seizure or control period) at each time segment, and repeated the same segment clusterization procedure to obtain 1000 surrogate statistic values. These values were used to approximate a null distribution against which we compared the original aggregated effect size value via a right-tail sided significance test. If the test’s significance value was below 0.05, we considered the pre-seizure interval formed by the adjacent segments to exhibit significantly lower centrality entropy than the one obtained in the control period and we identified it as a *critical phase*. In addition, we made use of the Kolmogorov Smirnov test to assess that the critical phase distribution across patients was significantly different from a distribution of randomly placed significant clusters of the same duration.

In general, to test paired or unpaired samples across time (e.g., pre-seizure vs. control period) or recording sites (e.g., seizure-onset sites across different periods) per patient, we made use of the Wilcoxon test for small sample sizes and the t-test for sufficiently large number of samples (>20). However, in most comparisons, non-comparable or very large number of samples could overestimate statistical effects. Hence, in those cases we computed and reported the effect size using Cohen’s d (based on the difference between medians/means for Wilcoxon test/t-test). To deal with the multiple-comparison problem, we applied the Holm-Bonferroni correction (Holm 1979) over patients in Fig 4 and over combination of regional comparisons in Fig 6. We resorted to linear regression and the coefficient of determination (R square) to evaluate the association between state probabilities/homogeneities and the decrease in centrality entropy. Finally, mean connectivity values across electrode pairs were computed using the Fisher transform (Fisher, 1920).

### Note on the typology of statistical tests

The main results combined within-subject and group-level statistical tests depending on the question at hand. Within-subject tests can be found in Fig 3A, Fig 4C (right) and Fig 6. Group-level tests can be found in Fig 3B and 3C, Fig 4C (left), and Fig 5B, 5C, 5D.

## Acknowledgements

We would like to thank Ralph Andrzejak, Maria Victoria Puig and Thomas Gener for their insightful comments during the preparation of this manuscript.

## Author contributions

A.T.C. and A.P. acquired the data; A.T.C. analysed the data and wrote the manuscript. All authors contributed to the design of the study paradigm and the interpretation of the results.

